# Correlation Analysis of Demographic Characteristics of Internal Medicine Residents on the Training Effect of Flipped Teaching Based on Video Conference

**DOI:** 10.1101/2022.08.13.503831

**Authors:** Xiao-Yu Zhang

## Abstract

**Objective:** I aimed to clarify whether the demographic characteristics of internal medicine residents are related to the training effect of Flipped Teaching based on Video Conference, so as to further improve the teaching management and quality control.

**Methods:** A total of 43 residents who participated in the Flipped Teaching based on Video Conference in April and June were included as the research objects. Online examination was conducted for these residents, and the demographic characteristics of these residents and their examination score were quantitatively analyzed, so as to clarify the correlation between the demographic characteristics of these residents and their examination score.

**Results:** The average score of 43 residents in the online examination was 72.0(64.0-79.0). 40 residents passed the exam, accounting for 93.0%. The bivariate correlations analysis showed that the age, gender, educational background, training batch and training phase of residents were not correlated with the examination score; all P > 0.05. The age, gender, educational background, training batch and training phase of these residents were not correlated with the examination score by single linear regression and multiple linear regression analysis; all P > 0.05.

**Conclusion:** Age, gender, educational background, training batches and training phases of internal medicine residents are not related to the examination score after Flipped Teaching based on Video Conference. Demographic characteristics may not be related to the training effect of residents through preliminary study.

## Introduction

In view of the fact that some standardized training bases for internal medicine residents did not meet the training standards of infectious diseases, those residents in the bases were assigned to a designated hospital to carry out the training of infectious diseases.[1] Due to the COVID-19 pandemic, Shanghai had implemented regional control measures since March 28, 2022[2]; Many internal medicine residents were unable to conduct infectious diseases training across the hospitals in April.

To continue the infectious disease training for internal medicine residents in Shanghai and ensure the quality of the training, I explored a new teaching model to carry out this training. From April 1 to April 4, I carried out Flipped Teaching based on Video Conference for those internal medicine residents scheduled to attend infectious disease training at designated hospitals in April, and this teaching model was generally effective in carrying out for internal medicine residents participating in the infectious diseases training, and it could be used as a supplementary training method for standardized training of internal medicine residents to make up for the shortage of actual training period in a certain stage;[3] So, I continued this training model throughout the whole April for infectious diseases training for those residents. For further verifying the training effect and repeatability of Flipped Teaching with Video Conference as carrier in the training of infectious diseases, I once more carried out this teaching model for those residents scheduled to attend infectious diseases training at the designated hospital in June, and the study showed that this teaching model was generally effective for internal medicine residents to carry out training of infectious diseases, with good feedback and strong feasibility.[4]

According to the requirements of standardized training for residents, it was necessary to conduct examination of residents training at the corresponding stage, and also further evaluate the training effect of this teaching model. I was conducting an online infectious disease examination for those residents on July 11. Through this examination, the effect and correlation of the demographic characteristics of those residents on the training effect of Flipped Teaching based on Video Conference was clarified, so as to further optimize the teaching management mode and quality control; Which could provide experience support for other organizations or institutions to adopt the teaching model.

## Objects and Methods

### Subjects

The study objects were internal medicine residents in April and residents in June who participated in the infectious diseases training with Flipped Teaching based on Video Conference. And they were also residents who had been participating in the standardized training in Shanghai.

Participant consents of those residents were obtained, including their data being used for the training and the research, and that this study was conducted in accordance with the Declaration of Helsinki.

### Organization and Implementation of Examination

The examination questions came from the standardized training question bank of internal medicine residents (infectious diseases), and were selected by the question bank software according to the classification of diseases. The online examination mode was adopted in this examination. Those residents needed to prepare two sets of equipment, one for online answering questions, and the other one for video surveillance. I jointly supervised the examination online with the teaching administrative personnel of the dispatched hospitals.

### Evaluation Indicators

Combined with the actual situation, this study included five indicators of age, gender, education background, training batch and training phase into demographic characteristics for statistical analysis.

### Software Application

The Questionnaire Star software was used to conduct online examination for those residents. The Tencent Conference software was used for online examination invigilation.

SPSS software version 23.0 (SPSS Inc. Chicago, IL, USA) was used for statistical analysis of the data. Continuous variables were expressed as medians and interquartile ranges to reflect the distribution of the study indicators. Categorical variables were represented by example (%) to reflect the composition ratio of the study indicators. Correlations between demographic characteristics indicators and examination score were analyzed by bivariate correlations, single linear regression and multiple linear regression. A P value of two-sided less than 0.05 was considered as statistically significant.

## Results

### Basic Information of Residents Participating in the Study

A total of 43 residents participated in the infectious diseases training with Flipped Teaching based on Video Conference, all from tertiary hospitals. Among them: 15 residents were males, accounting for 34.9%; average age was 29.0 years; 16 residents had bachelor degree, accounting for 37.2%; 8 residents had master degree, accounting for 18.6%; 19 residents had doctoral degree, accounting for 44.2%; 2 residents were in the first year of training, accounting for 4.7%; 29 residents were in the second year of training, accounting for 67.4%; 12 residents were in the third year of training, accounting for 27.9%; First batch had 19 residents in April, accounting for 44.2%; second batch had 24 residents in June, accounting for 55.8%; Average score of the examination was 72; And 40 residents passed the examination. The detailed information is shown in *Table 1*.

**Table 1.** Baseline of Residents Participating in Flipped Teaching

| Indicators |  |
| --- | --- |
| Total number [n, (%)] | 43 (100) |
| Sexuality [male, n, (%)] | 15 (34.9) |
| Age [years, median, (interquartile range)] | 29.0 (26.0–31.0) |
| Education background |  |
| Bachelor degree [n, (%)] | 16 (37.2) |
| Master degree [n, (%)] | 8 (18.6) |
| Doctor degree [n, (%)] | 19 (44.2) |
| Training phase |  |
| First year of training [n, (%)] | 2 (4.7) |
| Second year of training [n, (%)] | 29 (67.4) |
| Third year of training [n, (%)] | 12 (27.9) |
| From tertiary hospitals [n, (%)] | 43 (100) |
| Training batch for infectious diseases training in the designated hospital |  |
| First batch in April [n, (%)] | 19 (44.2) |
| Second batch in June [n, (%)] | 24 (55.8) |
| Examination score [years, median, (interquartile range)] | 72.0 (64.0–79.0) |
| Number of examination score $\geq 60$ / Number of passing examination [n, (%)] | 40 (93.0) |

### Bivariate Correlation Analysis Between Demographic Characteristics and Examination Score

Through bivariate correlation analysis, examination score had no correlation with age, gender, education background, training batch and training phase, all P > 0.05. The detailed bivariate correlation data is shown in *Table 2*.

**Table 2.** Correlations between demographic characteristic and Examination Score

| **Variables** | ***r* value** | ***p* value** |
| --- | --- | --- |
| Training phase | 0.104 | 0.507 |
| Age | -0.234 | 0.131 |
| Training batch | 0.096 | 0.542 |
| Education background | -0.264 | 0.088 |
| Gender | -0.004 | 0.978 |

### Linear Regression Analysis Between Demographic Characteristics and Examination Score

Through unitary linear regression analysis, examination score was not correlated with age, gender, education background, training batch and training phase, all P > 0.05; Through multiple linear regression analysis, examination score was not correlated with age, gender, education background, training batch and training phase, all P > 0.05. The detailed linear regression data is shown in *Table 3*.

**Table 3.** Linear Regression Analysis for Demographic Characteristics on Examination Score

| Variables | Single linear regression |  | Multiple linear regression |  |
| --- | --- | --- | --- | --- |
|  | *B* value (95% CI) | *p* value | *B* value (95% CI) | *p* value |
| Training phase | 0.811 (−3.654–7.275) | 0.507 | 0.709 (−5.748–7.165) | 0.825 |
| Age | −0.701 (−1.618–0.217) | 0.131 | −0.274 (−2.153–1.605) | 0.769 |
| Training batch | 1.748 (−3.990–7.486) | 0.542 | 1.073 (−5.016–7.161) | 0.723 |
| Education background | −2.660 (−5.730–0.410) | 0.088 | −1.809 (−8.430–4.813) | 0.583 |
| Gender | −0.081 (−6.087–5.926) | 0.978 | 1.452 (−5.012–7.916) | 0.652 |

## Discussion

This research was based on the previous study of exploration and verification of Flipped Teaching with Video Conference as the carrier for internal medicine residents,[3,4] a deeper level of data mining; which preliminarily clarified whether the demographic characteristics of those residents were correlated with the effect of this teaching model, so as to provide a basis for improvement of training design and personnel management in the future. To this end, we carried on this study and following seven aspects of analysis and elaboration were shown for discussion to this study.

### Interpretation of Relevant Data in the Study

#### Age

The median age of the residents in the two batches was 29 years, and the quartiles range were from 26 to 31 years. The study showed that there was no correlation between age and examination score. The result reflected residents of different ages could be competent for this teaching model in a certain age range, without special restrictions.

#### Gender

The proportion of female in the two batches of those residents was high, which was 52.6%[3] and 75.0%[4] respectively, accounting for 65.1% in total; There was no clear difference in the examination score between female and male residents.

#### Education

This study confirmed that the bachelor, master and doctor degree of those residents had no correlation with the examination score; And this was reflected in the fact that the standardized training of residents did not require excessive attention to academic qualifications, and did not support the integration of residents standardized training and academic education.

#### Teaching batches

This study suggested that there was no correlation between the two batches of Flipped Teaching and examination score, which would be related to the same management mode of the two batches of Flipped Teaching. Moreover, many residents in the second batch were already in clinical practice, which suggested that the effect of this teaching model was relatively stable, and that Flipped Teaching could be synchronized with clinical practice.

#### Training phase

This study suggested that those residents had no relation with the examination score in the training phase of first year, second year, or third year. Therefore, residents training in the same clinical department would not need to be completed in several different phrase.

### Comparison with Related Literature

*Flipped Teaching* was added in the *All Fields* of the query box from PubMed Advanced Search Builder and a total of 1212 records are displayed;[5] *Flipped Classroom* was added in the *All Fields* of the query box from PubMed Advanced Search Builder and a total of 970 records are displayed, as of 15:15 pm CET, August 9, 2022.[6] All the above reflected that research on Flipped Teaching or Flipped Classroom attracted more attention. With the publication of the paper “ Educators propose ‘flipping’ medical training” in 2012[7], Flipped Teaching had been studied and applied more and more in the field of medical education. Khe Foon Hew et al suggested that the flipped classroom approach in health professions education yields a significant improvement in student learning compared with traditional teaching methods.[8] Robert Connor Chick et al proposed that flipped classroom model might help to bridge the educational gap for surgical residents during this unprecedented circumstance during the COVID-19 Pandemic.[9] R C Chick et al found that utilization of a flipped classroom method was well received and preferred by surgical trainees, and it increased performance on pre-conference quizzes without increasing preparation time.[10] Jeff Riddell et al found that the flipped classroom and lecture were essentially equivalent as the differences were small in the crossover study.[11] Susan M Martinelli et al suggested that flipped classroom approach to didactic education resulted in a small improvement in knowledge retention and was preferred by anesthesiology residents.[12] Cliff Lee et al found that flipped classroom model was effective in educating students on periodontal periodontal diagnosis and treatment planning and was well received by the students.[13] Kelly L Graham et al found that flipped classroom during a 6-hour cardiovascular prevention curriculum showed greater effectiveness in knowledge gain compared with a standard approach in an ambulatory residency environment.[14] Antoine Marchalot et al suggested that blended learning (associating internet-based learning and flipped classroom) showed positive effective on the anaesthesia and critical care residents’ knowledge by increasing their homework’s time.[15] Sakina Sadiq Malik et al suggested that flipped classroom was found to be an effective teaching strategy for procedural skills in dermatology residents from the difference analysis between median pre-test and post-test scores in both groups.[16] Randy Y Lu et al suggested that flipped classroom method was received favorably by trainees and may complement traditional methods of teaching from the primary outcome measured residents’ preferences for classroom styles and the secondary outcome compared knowledge acquisition.[17] Fady Girgis et al suggested that flipped classroom for neurosurgery resident education was a viable approach to resident education and was associated with increased engagement and improved performance using validated knowledge-assessment tools.[18] Howard S Moskowitz et al suggested this type of curriculum, which combined a variety of approaches including a flipped classroom model with active participation and integrates app technology, could improve otolaryngology-head and neck surgery residents performance on educational assessments.[19] Eunicia Tan et al suggested that flipped classroom showed promise as an acceptable approach to in-house emergency medicine teaching.[20] All the above represented flipped teaching had achieved good feedback and application effect for residents training in the field of different clinical specialties. In my previous research, Flipped Teaching based on Video Conference was applied in the field of infectious disease training for internal medicine residents, which also achieved good feedback and effect;[3-4] Although there was no unified definition and implementation of Flipped Teaching in the teaching field so far, and Inconsistent background and application domain of flipped teaching, there were great differences in teaching practice. However, no research had been found on the relationship between demographic characteristics and residents training; My study first clarified the relationship between the both, and raised possible controversial issues in the field of standardized resident training; This study also reflected the importance and prospective of my study.

### Key Points of the Study

#### Examination questions sector

Although the reliability and validity of the examination questions were not tested, the examination questions came from the standardized training question bank of internal medicine residents (infectious diseases), and were randomly selected by the question bank software according to the classification of diseases; Therefore, this examination had certain reliability and validity.

#### Invigilation sector

Two sets of equipment were prepared for the examination online, one for online answering questions, and the other one for video surveillance. I jointly supervised the examination online with the teaching administrative personnel of the dispatched hospitals. so as to ensure the notary of the exam and the accuracy of the research data.

#### Management sector

Because the both teaching adopted vertical management model[21], the same personnel for organization and management; It could avoid the difference of both batches of teaching caused by artificial reasons, which might affect the reliability of statistical results.

### Advantages of the Study

#### Methodological sector

Three statistical methods were used to analyse the correlation between demographic characteristics and examination score to ensure the reliability of preliminary conclusions.

#### Research objects sector

Since all the research objects came from tertiary hospitals in the same region, the stability of the research samples was good.

### Limitations of the Study

Due to the small number of samples and the small proportion of failing the examination, logistic regression was not suitable to analyze the influencing factors of the teaching examination score.

This study examination to those residents did not conduct a self-comparison to compare the difference in examination score before and after the Flipped Teaching based on Video Conference.

### Subsequent Research Work

In order to further verify the correlation between demographic characteristics of residents and training effect of this teaching model, more samples were needed to conduct the study. Alternatively, other scholars with a large sample size could also verify the accuracy of the previous correlation between the both.

All the indicators in this study showed negative results. In the future, the screening of possible influencing indicators could be expanded accordingly, and other related factors or influencing factors can be identified, so as to optimize the training program and improve the teaching quality.

### Key Issues to Be Discussed

Some administrative regions combined standardized training of residents with academic education; However, the preliminary result of my study suggested that education background and training results were irrelevant, and it did not support the integration of standardized training for residents and academic education.

As the primary stage of the medical profession, the standardized training of residents mainly focuses on the training of clinical knowledge and clinical skills. Does it need to be integrated standardized training for residents with academic education at the residents level? Can residents standardized training be carried out simultaneously with academic education? These are clear views that need to be confirmed by more scholars with more data.

## Conclusions

There is no correlation between the effect of Flipped Teaching and the demographic characteristics of residents such as age, gender, educational background, training batches and training stages. For residents with these characteristics, there is no need to adjust the teaching plan to achieve smooth training or available results of this teaching model.

## Ethical Approval and Consent to Participate

Informed consents of participants in the standardized training for those residents were obtained for the training and the study. The study received Institutional Review Board (IRB) approval by the Shanghai Public Health Clinical Center Ethics Committee. The IRB number was No. 2021-S026-01.

## Acknowledgments

This study was supported by the internal medicine residents who had participated in the standardized training for residents in Shanghai. Thanks to the teaching administration departments and the resident standardized training bases of the united training hospital for their supports.

## Authors’ Contributions

Xiao-Yu Zhang made conception, design, acquisition of data, analysis and interpretation of data, drafted and revised the manuscript, and agreed to be accountable for all aspects of the work.

## Funding

This research received no external funding.

## Disclosure

The author declares no competing financial and / or non-financial interests.

## Consent for publication

The author has read and agreed to the published version of the manuscript.

## Literature share statement

The study included in the manuscript submitted to the journal is transparent. Consent from the corresponding author is required for any institution or individual to reprint this document.

## Notes

### Competing Interest Statement

The authors have declared no competing interest.

